# Image processing approaches to enhance perivascular space visibility and quantification using MRI

**DOI:** 10.1101/609362

**Authors:** Farshid Sepehrband, Giuseppe Barisano, Nasim Sheikh-Bahaei, Ryan P Cabeen, Jeiran Choupan, Meng Law, Arthur W. Toga

**Author notes:** **Correspondence to:** Farshid Sepehrband, PhD, Laboratory of Neuro Imaging, USC Mark and Mary Stevens Neuroimaging and Informatics Institute, Keck School of Medicine of USC, University of Southern California, Los Angeles, CA, USA, E.

## Abstract

Imaging the perivascular spaces (PVS), also known as Virchow-Robin space, has significant clinical value, but there remains a need for neuroimaging techniques to improve mapping and quantification of the PVS. Current technique for PVS evaluation is a scoring system based on visual reading of visible PVS in regions of interest, and often limited to large caliber PVS. Enhancing the visibility of the PVS could support medical diagnosis and enable novel neuroscientific investigations. Increasing the MRI resolution is one approach to enhance the visibility of PVS but is limited by acquisition time and physical constraints. Alternatively, image processing approaches can be utilized to improve the contrast ratio between PVS and surrounding tissue. Here we combine T1- and T2-weighted images to enhance PVS contrast, intensifying the visibility of PVS. The Enhanced PVS Contrast (EPC) was achieved by combining T1- and T2-weighted images that were adaptively filtered to remove non-structured high-frequency spatial noise. EPC was evaluated on healthy young adults by presenting them to two expert readers and also through automated quantification. We found that EPC improves the conspicuity of the PVS and aid resolving a larger number of PVS. We also present a highly reliable automated PVS quantification approach, which was optimized using expert readings.

## Introduction

Imaging the perivascular spaces (PVS), also known as Virchow-Robin space, has significant clinical value. Many recent studies have shown pathological alteration of PVS in a range of neurological disorders (Bacyinski et al., 2017; Banerjee et al., 2017; Brown et al., 2018; Cavallari et al., 2018; Feldman et al., 2018; Kalaria, 2018; Krueger and Bechmann, 2010; Laveskog et al., 2018; Park et al., 2017). It is also believed that PVS plays a major role in the clearance system, accommodating the influx of CSF to brain parenchyma through peri-arterial space, and the efflux of interstitial fluid to the lymphatic system through peri-venous space (Rasmussen et al., 2018; Tarasoff-Conway et al., 2015; Bacyinski et al., 2017; Iliff et al., 2012). MRI is a powerful tool that enables *in vivo, non-invasive* imaging of this less-known glia-lymphatic pathway.

In the clinical practice, PVS is quantified based on the number of visible PVS on the axial slice of a T2-weighted (T2w) image that has the highest number of PVS in the region of interest (Wardlaw et al., 2013). This process can be laborious and error-prone, so efforts to improve efficiency and accuracy have been made by using a wide range of automatic or semi-automatic segmentation techniques, from classical image processing approaches to deep neural network modelling (Ballerini et al., 2018; Boespflug et al., 2018; Comulada, 2015; Del C. Valdes Hernandez et al., 2013; Descombes et al., 2004; Hou et al., 2017; Jung et al., 2018; Park et al., 2016; Ramirez et al., 2015; Wang et al., 2016; Zong et al., 2016). Park et al. used auto-context orientational information of the PVS for automatic segmentation (Park et al., 2016). Recently, Ballerini et al. showed that Frangi filtering (Frangi et al., 1998) could robustly segment PVS by extracting the vesselness map based on the PVS tubular morphology (Ballerini et al., 2018). More recently, Dubost et al. used convolutional neural network with a 3D kernel to automate the quantification of enlarged PVS (Dubost et al., 2019, 2018).

While these methods have improved the automated segmentation of PVS, less effort has been made to enhance the visibility of PVS through postprocessing means. A typical MRI session includes a variety of different sequences and utilizing different intensity profiles of these sequences can potentially improve PVS detection rate for both visual reading and automated segmentation. Combining MRI signal intensities has been used for other applications to achieve tissue-specific sensitivity. Van de Moortele et al. combined T1w and proton density images by dividing the latter by the former to improve signal non-uniformity at ultra-high field and optimize vessels visualization (Van de Moortele et al., 2009). MRI multi-modal ratio was also used to map cortical myelin content (Glasser and Van Essen, 2011; Rowley et al., 2015; Viviani et al., 2017) (for a systematic evaluation of the contrast enhancement via the combination of T1w and T2w see (Misaki et al., 2014)). Additionally, Wiggermann et al. combined T2w and FLAIR to improve the detection of multiple sclerosis lesions (Wiggermann et al., 2016).

Here we describe a multi-modal approach for enhancing the PVS visibility, which was achieved by combining T1w and T2w images that were adaptively filtered to remove non-structural high frequency spatial noise. Furthermore, we present an automated PVS quantification technique, which can be applied to T1w, T2w or the enhanced contrast. The efficacy of the Enhanced PVS Contrast (EPC) on both visual and automated detection is assessed and the reliability of the automated technique is examined.

## Method

Four folds of evaluations were performed. First, the visibility of PVS in EPC was qualitatively and quantitively examined. Second, the visibility of PVS to expert readers was evaluated by comparing the number of PVS counted in EPC and T2w images. Third, PVS automatic counting was introduced and evaluated. Fourth, the reliability of the PVS automated quantification was assessed using scan-rescan MRI data.

### MRI data

T1w and T2w images of the human connectome project (HCP) (Essen et al., 2013) were used in the analysis. Structural images were acquired at 0.7 mm^3^ resolution images. We used data from “S900 release”, which includes 900 healthy participants (age, 22–37 years). The preprocessed T1w and T2w images (Glasser et al., 2013; Milchenko and Marcus, 2013; Sotiropoulos et al., 2013) were used. In brief: the structural images were corrected for gradient nonlinearity, readout, and bias field; aligned to AC-PC “native” space and averaged when multiple runs were available; then registered to MNI 152 space using FSL (Jenkinson et al., 2012)’s FNIRT. Individual white and pial surfaces were then generated using the FreeSurfer software (Fischl, 2012) and the HCP pipelines (Glasser et al., 2013; Sotiropoulos et al., 2013). Among HCP subjects, 45 were scanned twice with scan-rescan interval of 139 ± 69 days; these scans were used to assess the reliability of the automated PVS quantification.

### Enhanced PVS Contrast (EPC)

**Figure 1** summarizes the steps required to obtain EPC. After preprocessing the data, T1w and T2w images were filtered using adaptive non-local mean filtering technique (Manjón et al., 2010). The Rician noise of the MRI images, calculated using robust noise estimation technique presented by Wiest-Daessle et al. (Wiest-Daesslé et al., 2008), was used as the noise level for non-local filtering (Manjón et al., 2010). To preserve PVS voxels while removing the noise, filtering was applied only on high frequency spatial noises. This was achieved by using a filtering patch with a radius of 1 voxel, which removes the noise at a single-voxel level and preserves signal intensities that are spatially repeated (Manjón et al., 2010). Finally, EPC was obtained by dividing filtered images (i.e. T1w/T2w).

**Figure 1.**
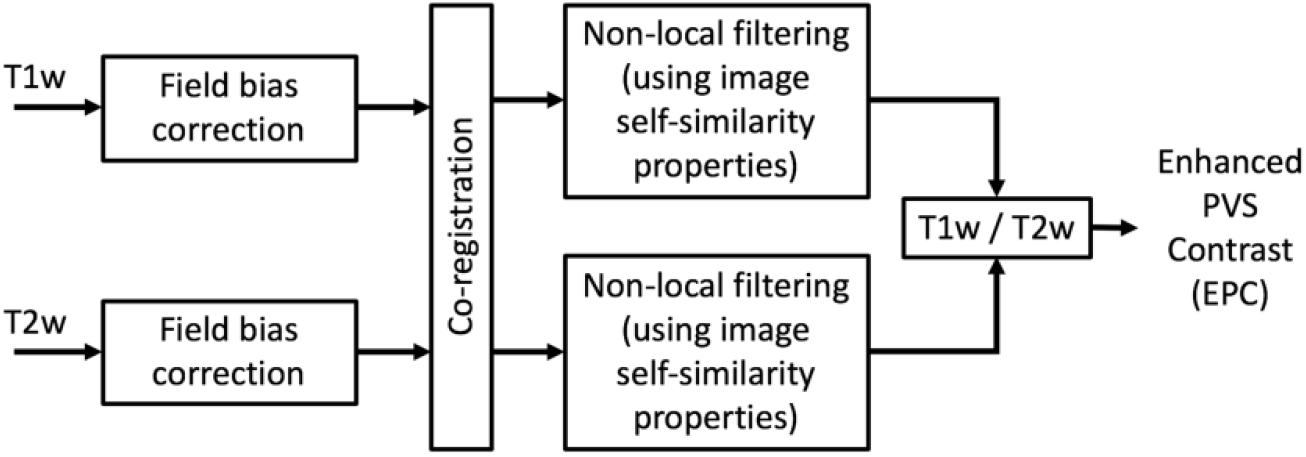
Flowchart of the Enhanced Perivascular space Contrast (EPC) technique.

PVS visibility was qualitatively compared across T1w, T2w, and EPC images in white matter and basal ganglia. PVS conspicuity was also assessed by comparing the PVS-to-white matter ratio in EPC images with that in T2, which was shown to provide a higher PVS contrast compared to T1w (Zong et al., 2016). A number of PVS and non-PVS white matter voxels were randomly selected across 10 subjects (more than 50 voxels per subject) and the average PVS-to-white matter ratio was measured.

### Expert reading and clinical scoring

PVS were independently rated in 100 randomly selected subjects of HCP by two expert readers (GB and NSB) on axial T2w and EPC images using a validated 5-point visual rating scale (Potter et al., 2015) in basal ganglia and centrum semi-ovale (0: no PVS, 1: 1-10, 2: 11–20, 3: 21–40, and 4: >40 PVS). Readers were blind to the rating results of each other. For validating the automated technique, the readers reached a consensus in few cases with different scores. One subject in which the PVS number was scored as “>100” by one of the readers was excluded from the statistical analysis. The total PVS score for each subject was calculated as the sum of the basal ganglia and centrum semi-ovale scores. The correlation between the number of PVS counted in T2w and EPC was calculated using Pearson correlation coefficient. PVS total counts were also compared using paired t-test. Lin’s concordance correlation (Lin, 1989) was used to determine the concordance between the two raters. In addition, inter-class correlation (ICC) estimates and their 95% confident intervals were calculated based on a mean-rating (k = 2), absolute-agreement, two-way mixed-effects model, as recommended in (Koo and Li, 2016). The statistical analysis was performed using SciPy library (version 1.2.0) on Python 3 and MATLAB statistics and machine learning toolbox.

### Automatic PVS quantification

We constructed a framework for PVS quantification and mapping using MRI, which can be applied to T1w, T2w, and EPC. Preprocessed data of HCP (Glasser et al., 2013) was used, which includes motion correction, non-uniform intensity normalization, Talairach transform computation, intensity normalization and skull stripping (Dale et al., 1999; Desikan et al., 2006; Fischl et al., 2004b, 2004a, 2002, 1999; Fischl and Dale, 2000; Reuter et al., 2012, 2010; Reuter and Fischl, 2011; Segonne et al., 2007, 2004; Sled et al., 1998; Waters et al., 2018). Then non-local mean filtering was applied and MRI images were parcellated to extract masks of white matter and basal ganglia, using *n*-tissue parcellation technique of the Advanced Normalization Tools (ANTs) software package (Avants et al., 2011). Parcellated brain (including white matter and basal ganglia) was used as a mask for PVS quantification analysis.

Subsequently, we applied Frangi filter (Frangi et al., 1998) to T1w, T2w, and EPC images using Quantitative Imaging Toolkit (Cabeen et al., 2018), which was implemented similar to (Ballerini et al., 2018). Frangi filter estimates a vesselness measure for each voxel 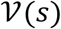 from eigenvectors *λ* of the Hessian matrix 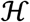 of the image:

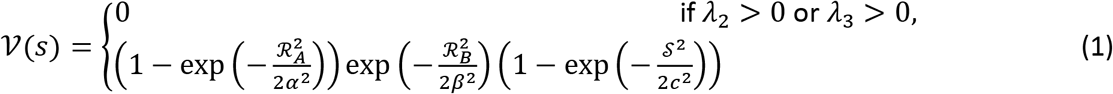

Where,

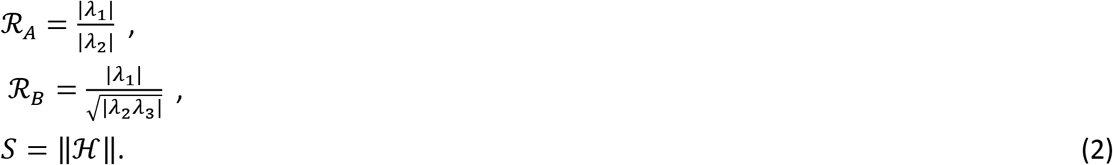

Default parameters of *α* = 0.5, *β* = 0.5 were used, as recommended in (Frangi et al., 1998). Frangi filter estimated vesselness measures at different scales and provided the maximum likeliness. The scale was set to a large range of 0.1 to 5 voxels in order to maximize the vessel inclusion. The output of this step is a quantitative map of vesselness in regions of interest, including white matter and basal ganglia.

In order to obtain a binary mask of PVS regions, the vesselness map should be thresholded. The binary mask enables automated PVS counting, volumetric, and spatial distribution analysis. Given that the vesselness value could vary across modalities, the threshold was optimized for each input image separately. We used the number of PVS counted by the experts for threshold optimization because of the absence of a ground truth. Vesselness values were standardized using robust scaling, in which values were scaled according to the inter-quartile range (IQR) to avoid the influence of large outliers:

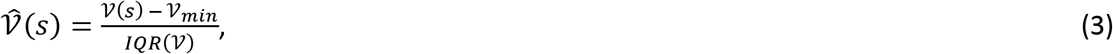

Then, the binary image of PVS mask was obtained by thresholding 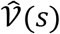.

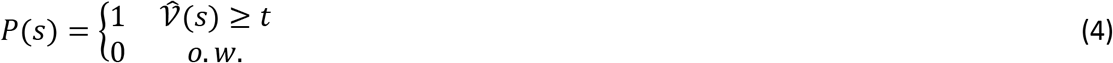

Then, the automated estimate of the total number of PVS was obtained by counting the number of connected components of the masked image *P*(*s*). Optimum thresholds were found by maximizing the concordance with the expert visual reading (concordance was used instead of absolute difference to optimize for global threshold and avoid biasing the threshold toward raters counting):

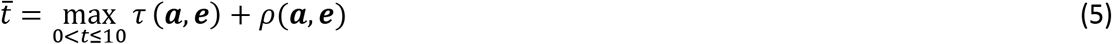

where ***a*** and ***e*** are one-dimensional arrays of PVS counts across all subjects (*n* = 100), obtained from the automated (*a*) and expert (*e*) readings, respectively. Kandall’s tau (*τ*) and Spearman’s Rho (*ρ*) were used to measure concordance and correlation, respectively. Average of expert readings from EPC images were used for optimization. Throughout the manuscript, the optimum thresholds of 2.3, 2.7 and 1.5 were used for T1w, T2w, and EPC, respectively.

After visual inspection, we noted that the imperfection of the white matter parcellation in periventricular and superficial white matter areas led to incorrected or missed PVS segmentation in white matter boundaries (an example is shown in **Supplementary Figure 1**). Therefore, the subtraction of a dilated mask of ventricles from the PVS mask was applied to exclude the periventricular voxels and remove the incorrectly segmented PVS at the lateral ventricles-white matter boundary. After obtaining the final PVS mask, the number of PVS was obtained by counting the number of connected components of the PVS mask. Small components (<5 voxels) were excluded from automated counting to minimize noise contribution. The automated technique was applied on all subjects. Finally, one-way ANOVA was conducted to compare the effect of input image (T1w, T2w, and EPC) on the estimated number of PVS.

### Test-retest reliability

We also evaluated the test-retest reliability of the PVS quantification with MRI. Forty MRI data from HCP (Essen et al., 2013) that included scan-rescan were used for reliability analysis. EPC was derived, and the automated quantification pipeline was applied on the scan-rescan images (T1w, T2w and EPC images): identical parameters and threshold were applied on scan-rescan data. Then scan-rescan reliability was assed using ICC, Lin’s concordance (Lin, 1989) and Pearson correlation analysis. ICC estimates and their 95% confident intervals were calculated based on a mean-rating (k = 2), absolute-agreement, two-way mixed-effects model.

## Results

### Evaluation 1: comparing the EPC with T1w and T2w

The PVS were more visible in EPC compared to T1w and T2w images (**Figures 2** and 3, and **Supplementary Figure 2**). The superiority of the EPC was evident in both white matter (**Figure 2**) and basal ganglia (**Figure 3**). EPC allowed the detection of PVS that were hardly identifiable in T1w and T2w (see yellow arrows in **Figure 2**: PVS that could barely be spotted in T1w and T2w were evident in EPC). PVS were more conspicuous in EPC compared to T1w and T2w. Moreover, PVS-to-white matter contrast was significantly higher in EPC compared to that in T2w (**Supplementary Figure 3**: ~2 times higher, *p*<0.0001).

**Figure 2.**
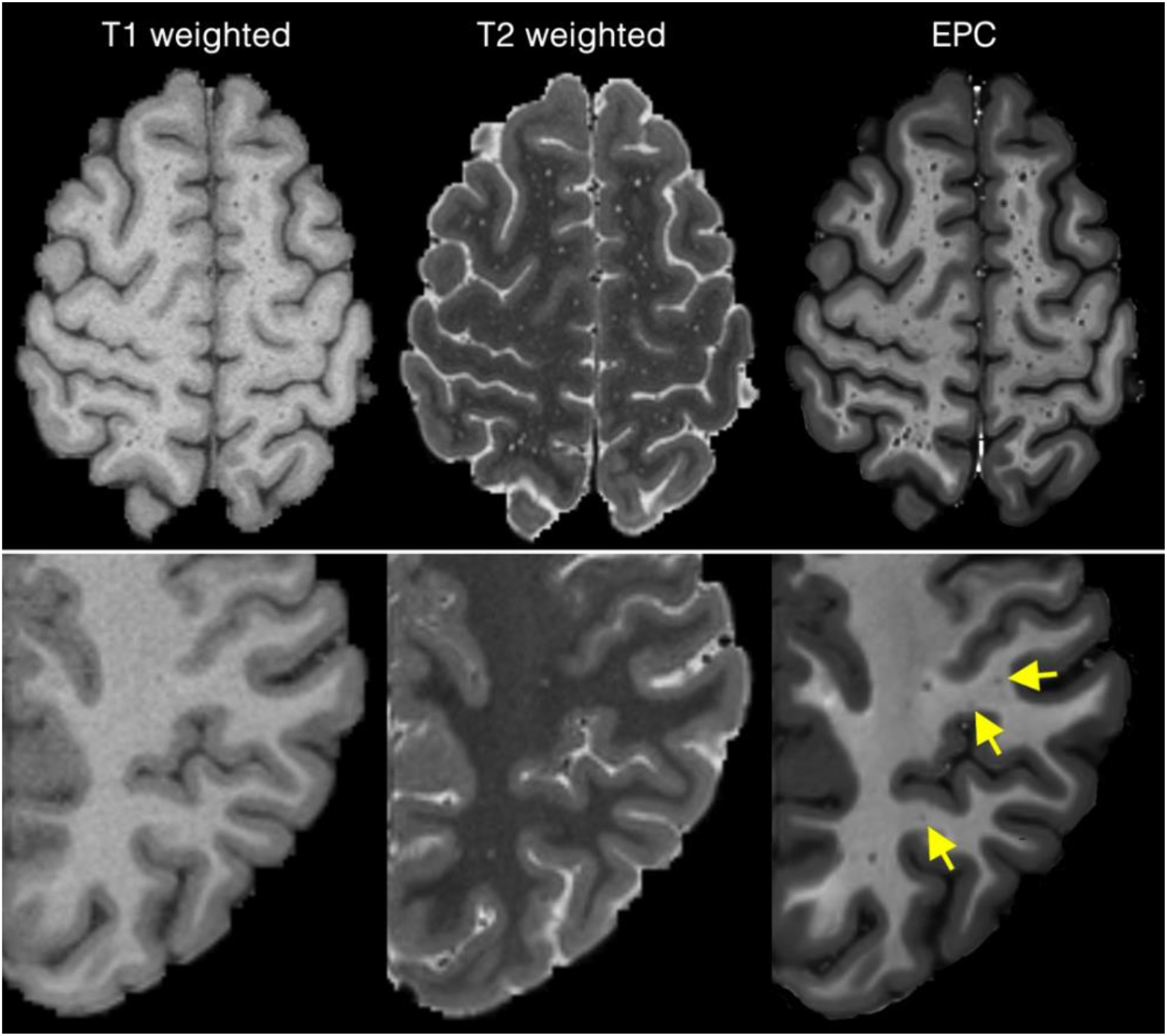
Comparing Enhanced PVS Contrast (EPC) with T1w and T2w images across two subjects with high perivascular spaces (PVS) presence (first row) and low PVS presence (second row).

**Figure 3.**
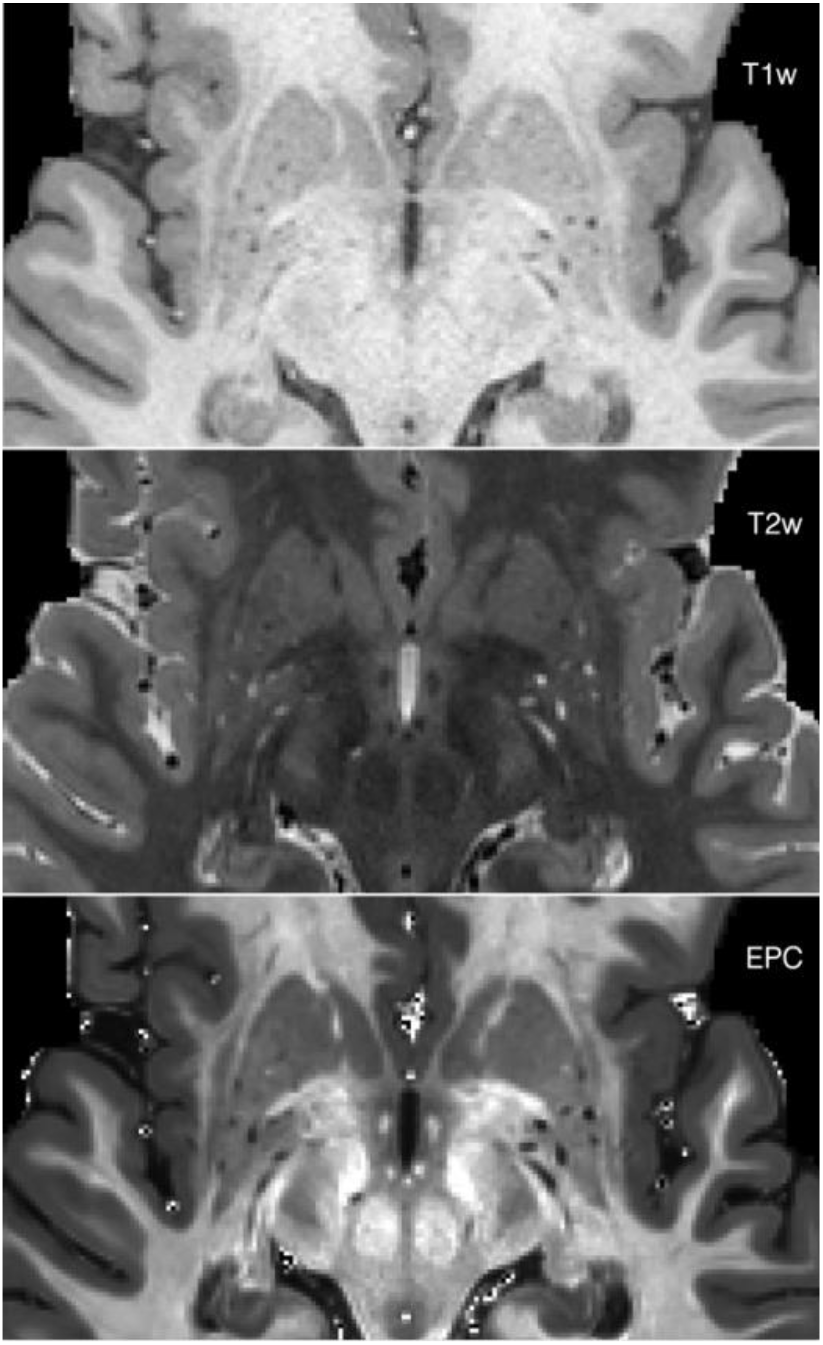
T1w, T2w, and the Enhanced PVS Contrast (EPC) images of the basal ganglia.

### Evaluation 2: Expert reads of PVS from EPC and T2w

A significantly higher number of PVS was counted in centrum semi-ovale by the readers when EPC was used compared with T2w (**Figure 4.a-b**: *t*(198)=5.8; *p*<0.0001 and **Figure 4.d-e**: *t*(198)=4.6; *p*<0.0001). PVS numbers measured in EPC and T2w were significantly correlated (*r*=0.81, *p*<0.0001 and *r*=0.8, *p*<0.0001 for the first and second expert, respectively). However, PVS individual categories were different: when EPC was used, the majority of the subjects were rated 4 (i.e. more than 40 PVS counted) and the rest were rated 3 (i.e. 21-40 PVS counted), whereas in T2w the majority of the subjects were rated 3. In fact, when T2w images were used, readers counted 34.4 ± 14.4 and 34.2 ± 15.2 PVS on average; in EPC, the average PVS counted by the experts increased to 46.5 ± 14.7 and 43.8 ± 14.1. High inter-rater reliability and concordance were observed, with a slight increase when EPC was used. The average ICCs of T2w and EPC were 0.96 and 0.98, with 95% confident intervals of 0.938-0.972 (*F*(99)=24.07, *p*<0.0001) and 0.966-0.985 (*F*(99)=43.69, *p*<0.0001), respectively. Lin’s concordance coefficients for T2w and EPC were 0.92 and 0.94, respectively.

**Figure 4.**
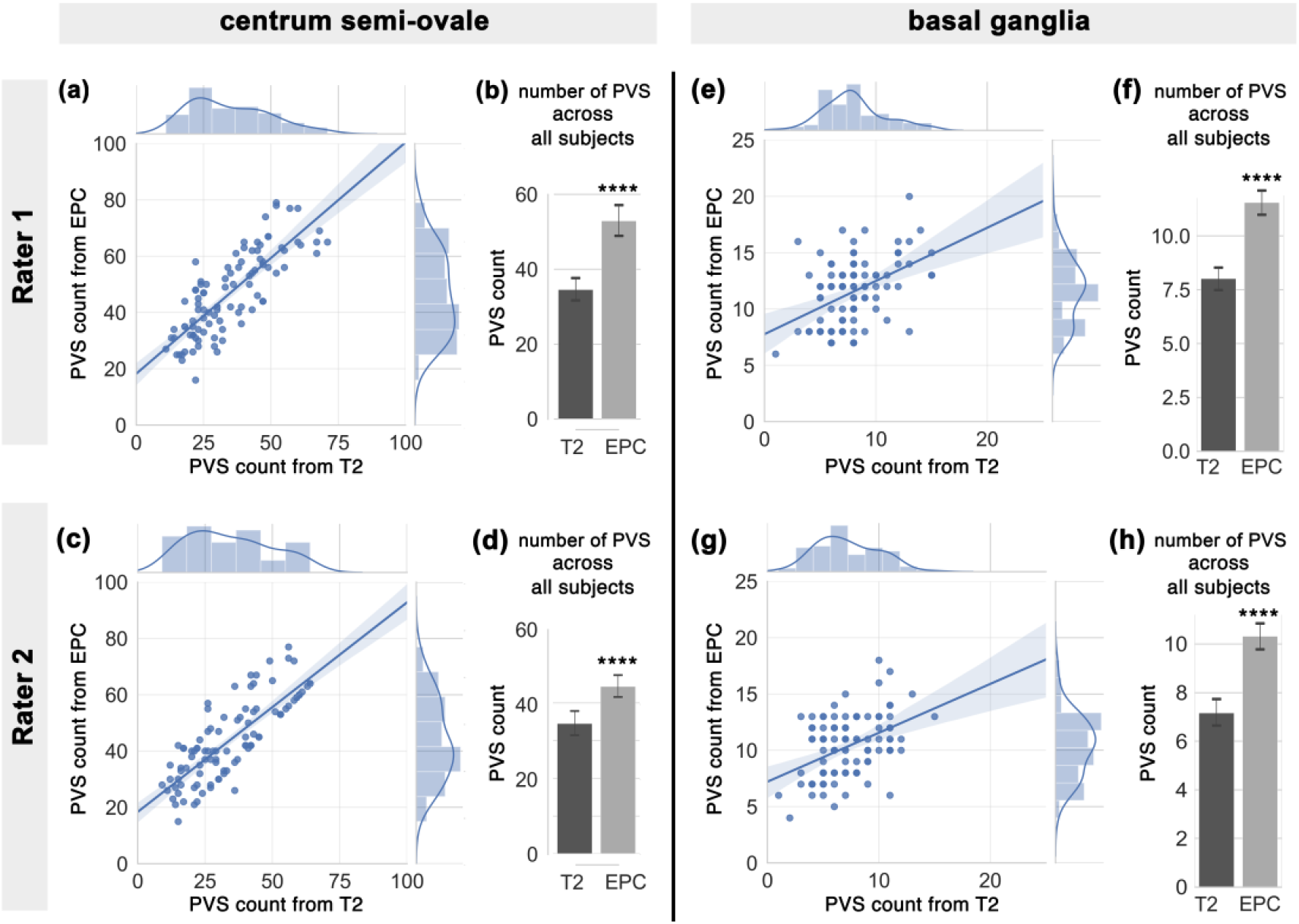
PVS number in centrum semi-ovale and basal ganglia across the analyzed subjects (N=99), counted in T2-weighted (T2w) and enhanced PVS contrast (EPC) images. **First row** refers to the first expert reader, and **second row** relates to the second expert reader. **(a)** and **(c)** plots show the correlation between the number of PVS counted on T2w and EPC images, and their distribution. **(b)** and **(d)** bars compare the mean and standard deviation of PVS across all subjects as derived from T2w and EPC images. **(e-h)** plots are similar to (a-d) but related to basal ganglia scores. Note that expert readers counted significantly higher number of PVS when EPC was used (*p*<0.0001).

A similar trend was observed in the basal ganglia (**Figure 4e-h**), where a significantly higher number of PVS were counted with EPC (t(198)=8.8; *p*<0.0001 and *t*(198)=8.14; *p*<0.0001, for the first and second reader, respectively). Compared to centrum semi-ovale results, a lower correlation between PVS number obtained from T2w and EPC was observed in basal ganglia (*r*=0.45, *p*<0.0001 and r=0.43, *p*<0.0001, for the first and second readers, respectively). When T2w images were used, readers counted 8.0 ± 2.8 and 7.2 ± 2.7 PVS on average; while the average PVS counts increased to 11.6 ± 2.9 and 10.3 ± 2.7 in EPC images. Furthermore, interrater reliability and concordance was slightly higher when EPC was used. The average ICCs of T2w and EPC were 0.87 and 0.92, with 95% confident intervals of 0.81-0.91 (*F*(99)=7.84, *p*<0.0001) and 0.87-0.94 (*F*(99)=11.86, *p*<0.0001), respectively. Lin’s concordance coefficients for T2w and EPC were 0.74 and 0.77, respectively.

### Evaluation 3: PVS automated quantification

A map of vessel likeliness overlaid on EPC is shown in **Figure 5**. PVS segmentation of the automated technique is dependent on the threshold applied. For EPC, 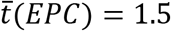 provided the optimum threshold which led to the highest concordance and Spearman’s correlation with expert readings. Qualitative inspection of the PVS masks obtained with different threshold highlights the superiority of the derived threshold in comparison to smaller or larger thresholds. Optimum threshold was different for different inputs: 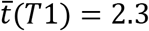 and 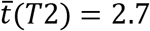 (**Figure 6**).

**Figure 5.**
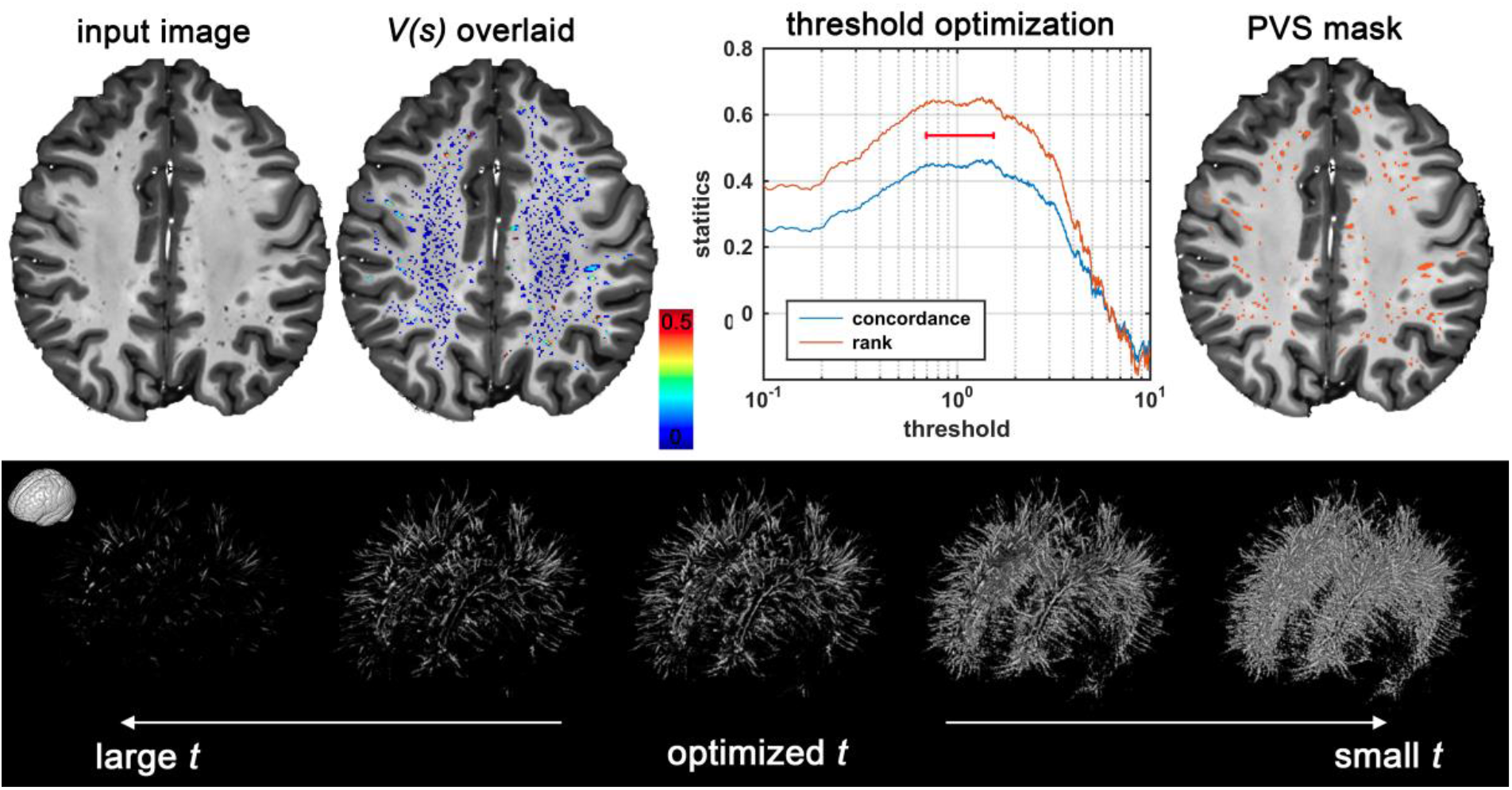
Automated quantification of PVS likeliness and PVS segmentation. First row shows an enhanced PVS contrast (EPC) image and the vesselness map, *V*(*s*), obtained from Frangi filtering (see method section and equation (1) for details). The optimization results to find an optimum global threshold for EPC is shown, in which *t* = 1.5 resulted to the highest concordance and correlation between automated PVS counting across the whole white matter and expert counting (number of PVS in the axial slice with the highest PVS presence). **Second row** highlights the influence of threshold selection on the PVS mask. A too small or too high threshold results in a large number of false positives or false negatives, respectively.

**Figure 6.**
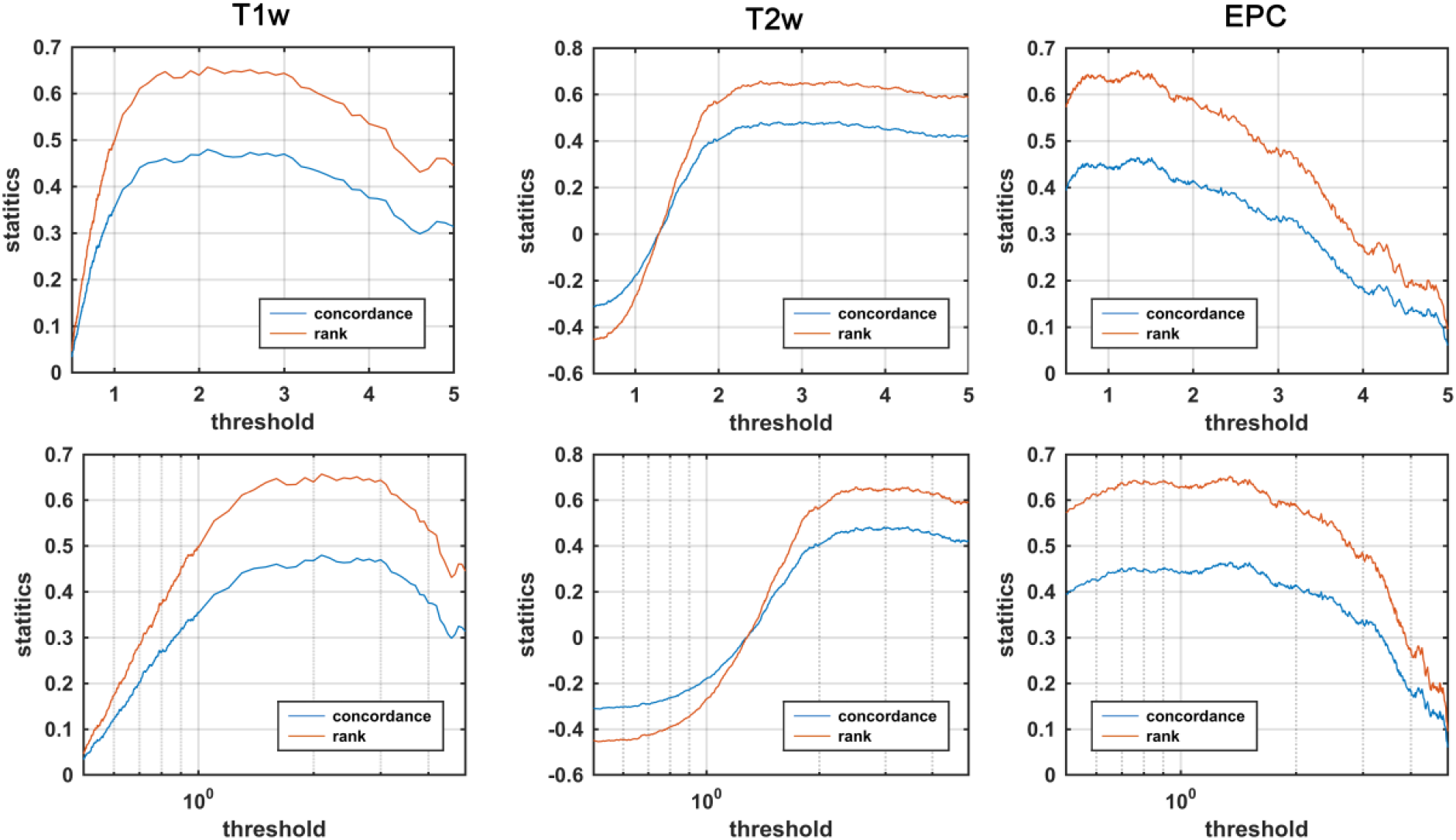
Threshold optimization for segmenting PVS for different modalities. Concordance and correlation of automated PVS quantification with expert readings are presented for different threshold values. Given that *threshold* appeared to be optimum within a relatively wide range, a visual inspection of the threshold is recommended.

Visual inspection showed that, as expected, the Frangi filter was able to detect the tubular structures of the PVS (**Figure 7**). No statistical evidence was found to suggest the automated quantification of PVS using EPC is superior to those derived from T1w and T2w. PVS quantification (number of PVS) were significantly correlated across T1w, T2w, and EPC results (all at *p*<0.0001) and all the automatic measurements reported a similar concordance level with the expert scores. Lin’s concordance coefficient between automated PVS counts and expert scores was 0.81, the bias correction value (a measure of accuracy) was 0.88, and the Pearson correlation coefficient was 0.91 (*p*<0.0001). The analysis of variance showed that the effect of input image (T1w, T2w, and EPC) on the number of PVS measured was significant *F*(2,294) = 48.56, *p* < 0.0001, which suggests that the automated technique for PVS quantification needs to be applied on the same image modality across study data.

**Figure 7.**
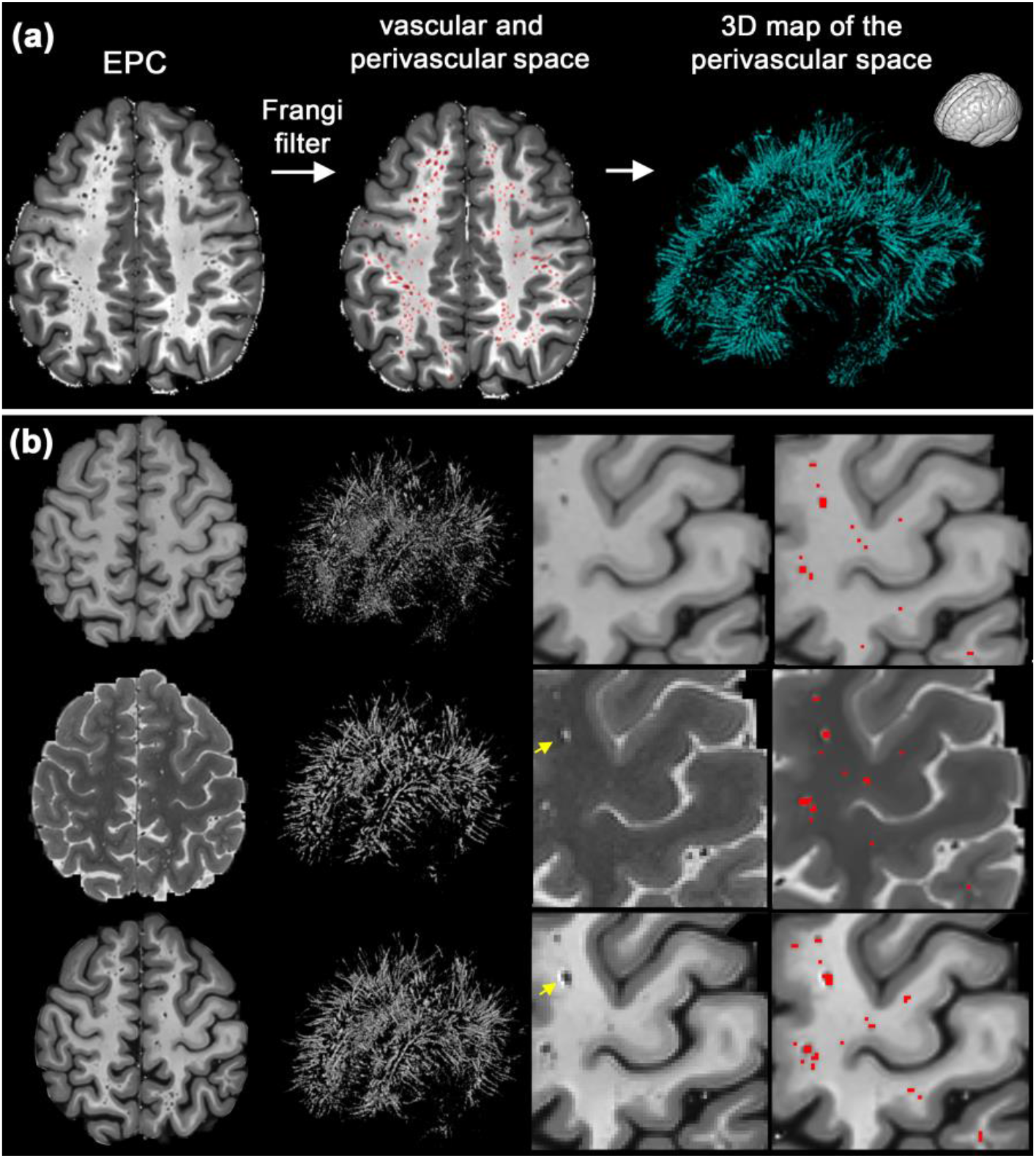
Mapping perivascular space (PVS). **(a)** an example of PVS segmentation using the Frangi filter with the optimum threshold. **(b)** Comparison of automated segmentation when T1w, T2w, and EPC were used as the input. **Supplementary Video 1** shows a 3D view of the PVS mask obtained from EPC. Yellow arrows show voxels with an artifact (which could be Gibbs ringing and/or internal gradient artifact). Given the inverse signature of this artifact in relation to PVS in both T2w and EPC, the PVS identification was unaffected.

### Evaluation 4: test-retest reliability of automated quantification

Excellent test-retest reliability was observed in PVS automated quantification regardless of the input image used (**Figure 8** and **Table 1**). Same thresholds, optimized on different subjects, were used for scan-rescan data. Lin’ concordance coefficient between scan-rescan PVS measurements were 0.89, 0.94 and 0.83 for T1w, T2w and EPC, respectively. T2w images showed the highest concordance compared to other inputs. PVS measures were significantly correlated between scan-rescan images (*r*=0.90, *p*<0.0001, *r*=0.95, *p*<0.0001, and r=0.85, *p*<0.0001 for T1w, T2w and EPC, respectively). For all three inputs, no difference between the number of PVS in scan and rescan was observed.

**Figure 8.**
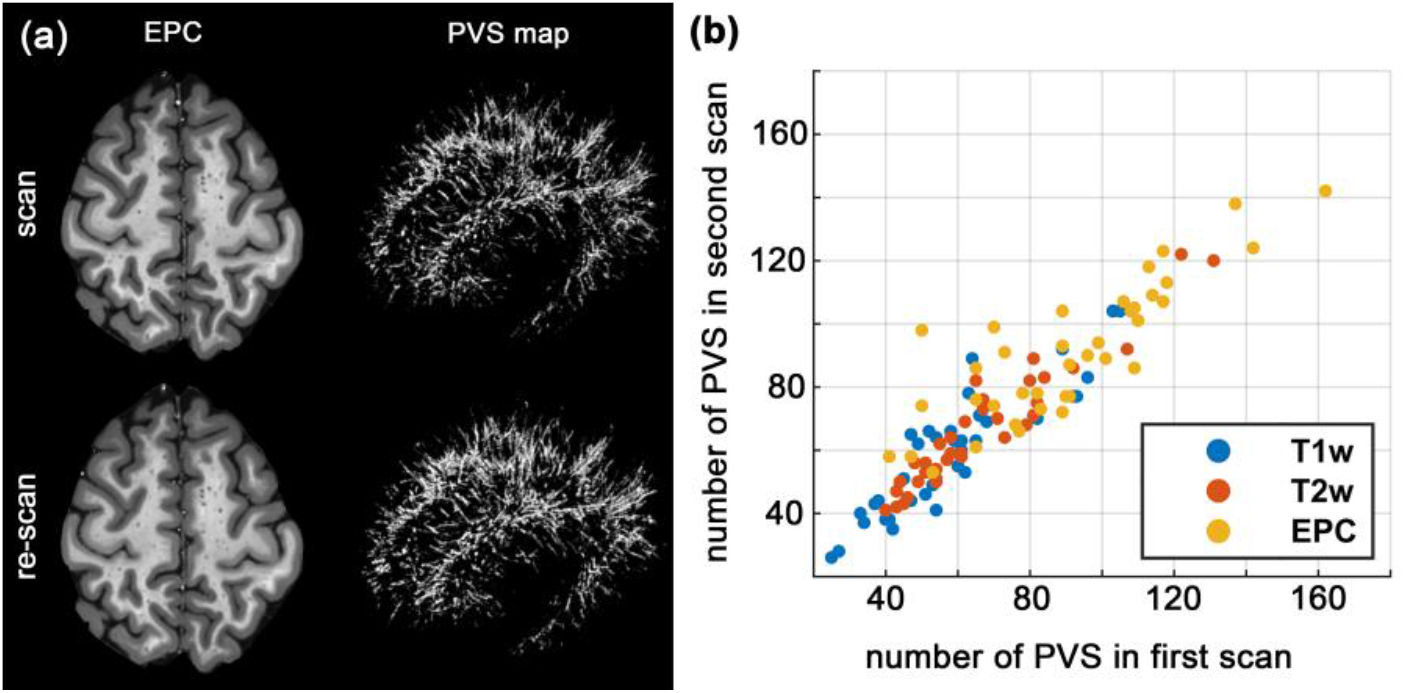
Automated PVS quantification reliability assessment using scan-rescan data. **(a)** An example of the scan-rescan maps of the enhanced PVS contrast (EPC) and the PVS mask obtained from the same subject. **(b)** The graph shows the total number of automatically counted PVS across subjects on the scan-rescan data, obtained from different inputs.

**Table 1.** Inter-class correlation (ICC) statistics of scan-rescan experiment for automated quantification of perivascular space (PVS), from T1w, T2w and enhanced PVS contrast (EPC) images.

| *Input image* | *ICC* | *95% CI*<br>*lower bound* | *95% CI*<br>*upper bound* | *F score* | *p-value* |
| --- | --- | --- | --- | --- | --- |
| **T1w** | 0.94 | 0.89 | 0.97 | 18.04 | <0.0001 |
| **T2w** | 0.97 | 0.95 | 0.99 | 38.49 | <0.0001 |
| **EPC** | 0.90 | 0.81 | 0.95 | 10.31 | <0.0001 |

## Discussion

In this article, we presented a combined T1w-T2w approach to enhance the visibility of the PVS. EPC was utilized for both expert and computer-aided readings and was evaluated qualitatively and quantitatively. The proposed map (EPC) enhances the contrast and improves the conspicuity of the PVS, resulting in detection of a significantly larger number of PVS identified by expert readers. EPC benefits from the inverse signal profile of fluid on T1w and T2w images: when these images are combined together, a magnified PVS-tissue contrast can be obtained. EPC also uses a spatial non-local mean filtering technique, which has shown to be effective for mapping PVS (Hou et al., 2017). We noted that even in the high-resolution T2w images of the human connectome project (0.7 mm^3^ resolution), it is difficult to detect small PVS (**Supplementary Figure 2**), while they could be identified with this new technique.

According to the current standard visual rating scale for PVS (Potter et al., 2015), most of the subjects in this cohort belonged to the class with the highest amount of PVS (category 4: >40 PVS). Moreover, the inter-subject variability of PVS was large (standard deviation of ~15), despite the fact that subjects are all young healthy adults. This highlights the fact that the current rating approach has inherent limitations, because 1) a counting approach is highly dependent on the image resolution and quality and is difficult to generalize, 2) dichotomizing the PVS count reduces the statistical power and could underestimate the extent of variation, because considerable variability may be absorbed within each group (Altman and Royston, 2006), 3) it does not consider the morphometric and spatial information of the PVS. In order to detect the intra- and inter-subject PVS alteration and ultimately to determine the role of PVS in different pathologies, there is a need to improve the current imaging and rating techniques.

With the advancement of the MRI technology, structural imaging in submillimeter resolution is achievable in a plausible time frame; for example, T1w MRI with 0.7mm^3^ resolution can be obtained in 8 minutes at 3T (Van Essen et al., 2012). Such imaging resolution enables to visualize PVS that were otherwise not apparent due to partial volume effect. With the added visibility of PVS comes the challenge of counting and mapping, as the visual rating becomes extremely laborious. Here we presented a pipeline that can be used to automatically map PVS.

The scan-rescan experiment showed that EPC is highly reliable, with no observed statistical difference across scan-rescan results. A trivial difference between scan-rescan PVS maps was observed, which are most likely due to **1)** segmentation imperfection and image intensity differences of scan-rescan signal (e.g. due to subject motion) (Madan and Kensinger, 2017; Reuter et al., 2015), **2)** normal physiological changes of PVS in the same subject, such as potential effects of time-of-day and hydration on morphometric estimates of PVS (Dickson et al., 2005; Kempton et al., 2009; Trefler et al., 2016).

Whether PVS quantification is done by a neuroradiologist or automatically, an image with high PVS-tissue contrast is ideal. Currently, the modality of choice for PVS analysis is T2w, because it offers a higher contrast of PVS-tissue compared to T1w (Zong et al., 2016). Yet, small PVS (< image resolution) are difficult to detect or separate from noise. Hence, PVS quantification is often limited only to enlarged PVS, despite the fact that physiological or pathological changes are expected to initiate in submillimeter scale (Horsburgh et al., 2018; Shi and Wardlaw, 2016). In order to improve the mapping of the PVS, the MRI contrast of the PVS and the neighboring parenchyma should be increased. Besides the image processing approaches, PVS contrast can be enhanced through MRI technological improvement such as optimizing imaging sequence (Zong et al., 2016) and employing ultra-high field technology (Barisano et al., 2018; Hou et al., 2017; Park et al., 2016; Zong et al., 2016).

Our quantitative PVS mapping and previous works showed that PVS can be mapped from an individual MRI contrast (Ballerini et al., 2018; Boespflug et al., 2018; Comulada, 2015; Del C. Valdes Hernandez et al., 2013; Descombes et al., 2004; Hou et al., 2017; Jung et al., 2018; Park et al., 2016; Ramirez et al., 2015; Wang et al., 2016; Zong et al., 2016). However, it should be noted that in the presence of pathology, additional image sequences (e.g. FLAIR) can be useful to discern PVS from pathological changes, such as white matter hyperintensities (WMH). Another advantage of a multi-modal approach for PVS quantification is the improvement of misclassification. For instance, vessels not surrounded by PVS are not easily distinguishable from vessels with PVS if only T1w is used, because both appear hypointense in this modality. T2w and EPC are able to solve this issue: in fact, in the absence of PVS, vessels appear hypointense in T2w and hyperintense in EPC, unlike vessels with PVS, which appear hyperintense in T2w and hypointense in EPC (see **Supplementary Figure 4**). Additionally, the correct identification of PVS can be achieved via the analysis of its morphometric characteristics, including size, shape, and anatomical location. Ballerini et al., for example, argued that Frangi filter ensures specificity in PVS segmentation given the tubular structure of the PVS.

Two recent studies have also used multi-modal techniques for PVS segmentation and showed that it outperformed segmentation derived from a single modality (Ballerini et al., 2018; Boespflug et al., 2018). Ballerini et al. applied segmentation on each modality and combined segmentations outputs using an AND operation. Boespflug et al. applied multivariate clustering, followed by morphometric filtering, to extract PVS from T1w, T2w, FLAIR and proton density images. It should be noted that the aim of these techniques was to improve the accuracy of the automated segmentation, but our study aimed to propose a map that improves the visibility and detectability of the PVS, which can also make the visual scoring more accurate.

In addition to its potential clinical relevance, mapping PVS can be useful to improve the accuracy of other quantitative MRI techniques as well, because these could be affected by the partial presence of PVS in image voxels. Recently, we have shown that ignoring PVS could systematically affect how diffusion tensor imaging measures can be interpreted (Sepehrband et al., 2019). Such contribution and its potential confounding effect may be ameliorated if PVS is mapped and included in the analysis.

A limitation of multi-modal combination techniques is that it requires additional scan time and therefore is more prone to subject motion, which could negatively affect the co-registration. An interleaved acquisition was shown to be highly effective to ameliorate the co-registration issue (Van de Moortele et al., 2009). Another limitation of EPC is that it requires the same image resolution for T1w and T2w. These sequences are often acquired in different resolutions, particularly in clinical practices (T2w are often acquired with thicker axial slices). Future investigations could focus on determining the extent to which the resolution difference between T1w and T2w affects the EPC quality and whether an intra-subject co-registration could amend this limitation. Finally, imperfection of the brain parcellation could affect the automated quantification (see an example in **Supplementary Figure 1**). Further efforts are required to explore the effect of brain parcellation on PVS mapping or to build computational tools that minimize the parcellation dependency.

## Conclusions

Our combined T1w-T2w approach (EPC) has demonstrated to enhance the visibility of the PVS, resulting in improvement of PVS mapping. EPC allowed both the expert readers and the computer-aided algorithm to achieve a more accurate and precise quantification of PVS. This novel method, which can be easily applied to a number of MRI datasets, aims to overcome the limitations of current MRI sequences in PVS detection and quantitative analysis. This is relevant not only to better characterize the role of PVS when they are enlarged in pathological conditions, but especially to perform quantitative research on PVS when they are small, such as in physiological and prodromal states.

## Acknowledgement

This work was supported by NIH grants: 2P41EB015922-21, 1P01AG052350-01 and USC ADRC 5P50AG005142. The content is solely the responsibility of the authors and does not necessarily represent the official views of the NIH.

**HCP:** Data were provided by the Human Connectome Project, WU-Minn Consortium (Principal Investigators: David Van Essen and Kamil Ugurbil; 1U54MH091657) funded by the 16 NIH Institutes and Centers that support the NIH Blueprint for Neuroscience Research; and by the McDonnell Center for Systems Neuroscience at Washington University.

## Supplementary figures

**Supplementary Figure 1.**
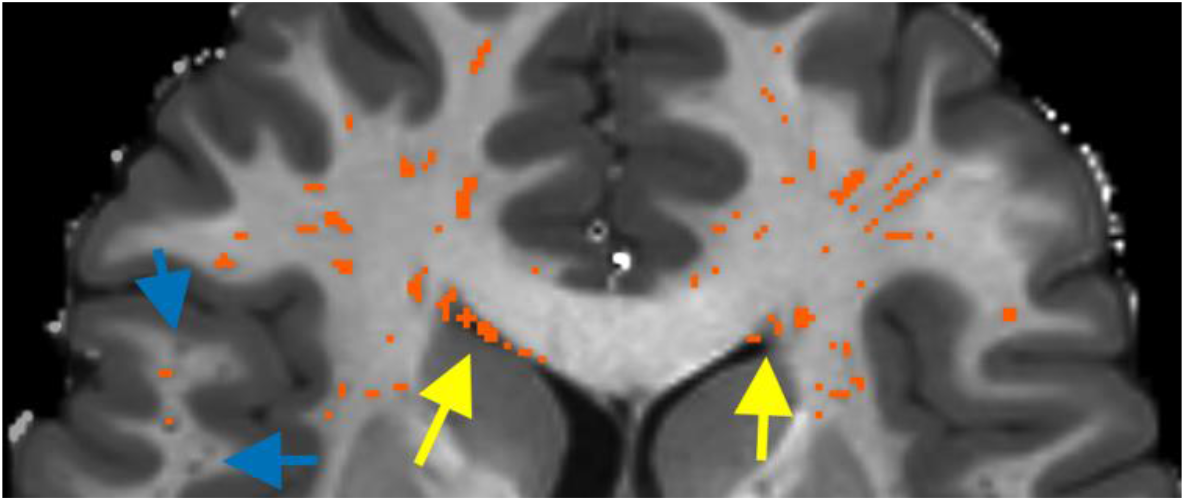
Examples of negative influence of the imperfect white matter parcellation on perivascular spaces (PVS) segmentation. Yellow arrows show periventricular voxels misclassified as PVS. Blue arrows show voxels with PVS in the superficial white matter which were not included in the white matter parcellation. While false positive voxels in periventricular area were removed by applying a dilated mask of the ventricle, the superficial white matter voxels remained unsolved.

**Supplementary Figure 2.**
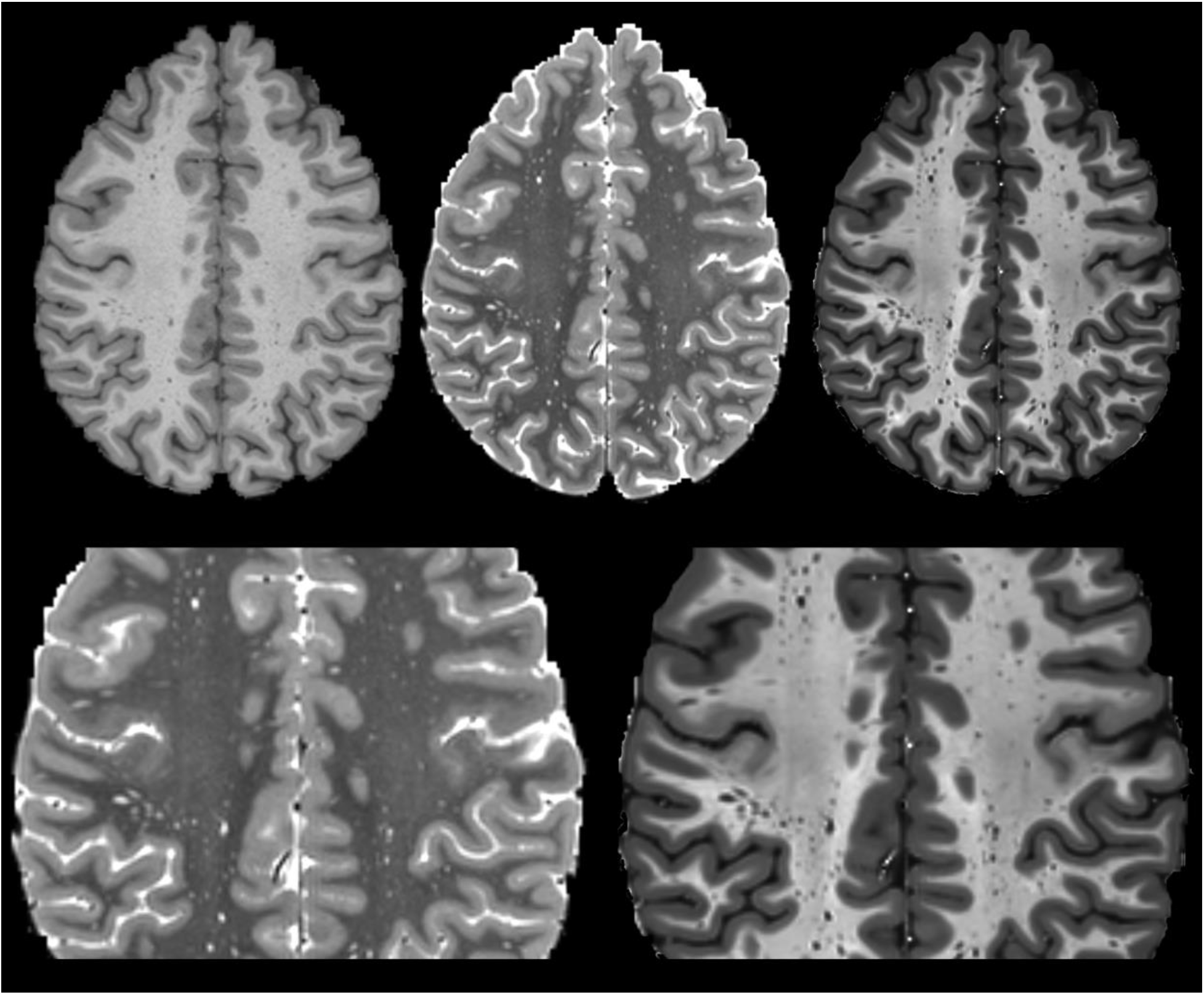
Comparing Enhanced PVS Contrast (EPC) with T1-weighted and T2-weighted images in subject with large number of small perivascular spaces (PVS) in centrum semi-ovale. Note that EPC aids visual identification of PVS and distinguishing them from image noise.

**Supplementary Figure 3.**
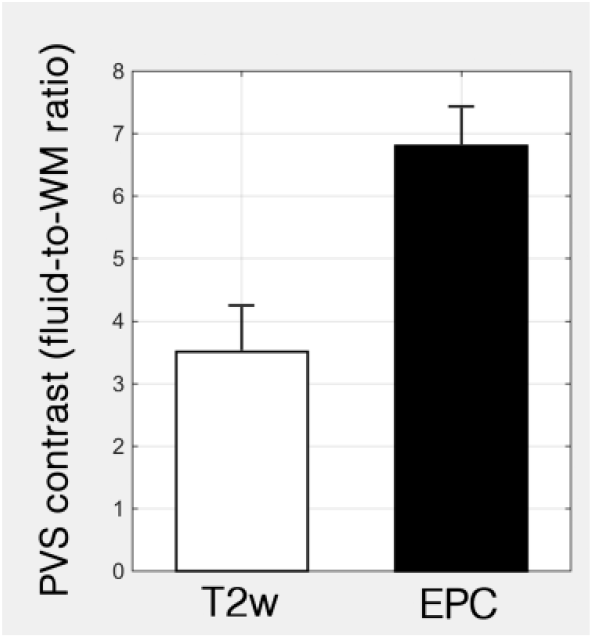
The contrast ratio between perivascular spaces (PVS) and the adjacent white matter voxels from multiple manually selected regions. The PVS-to-white matter contrast ratio of the EPC was significantly (*p*<0.0001) higher than that derived from T2w images.

**Supplementary Figure 4.**
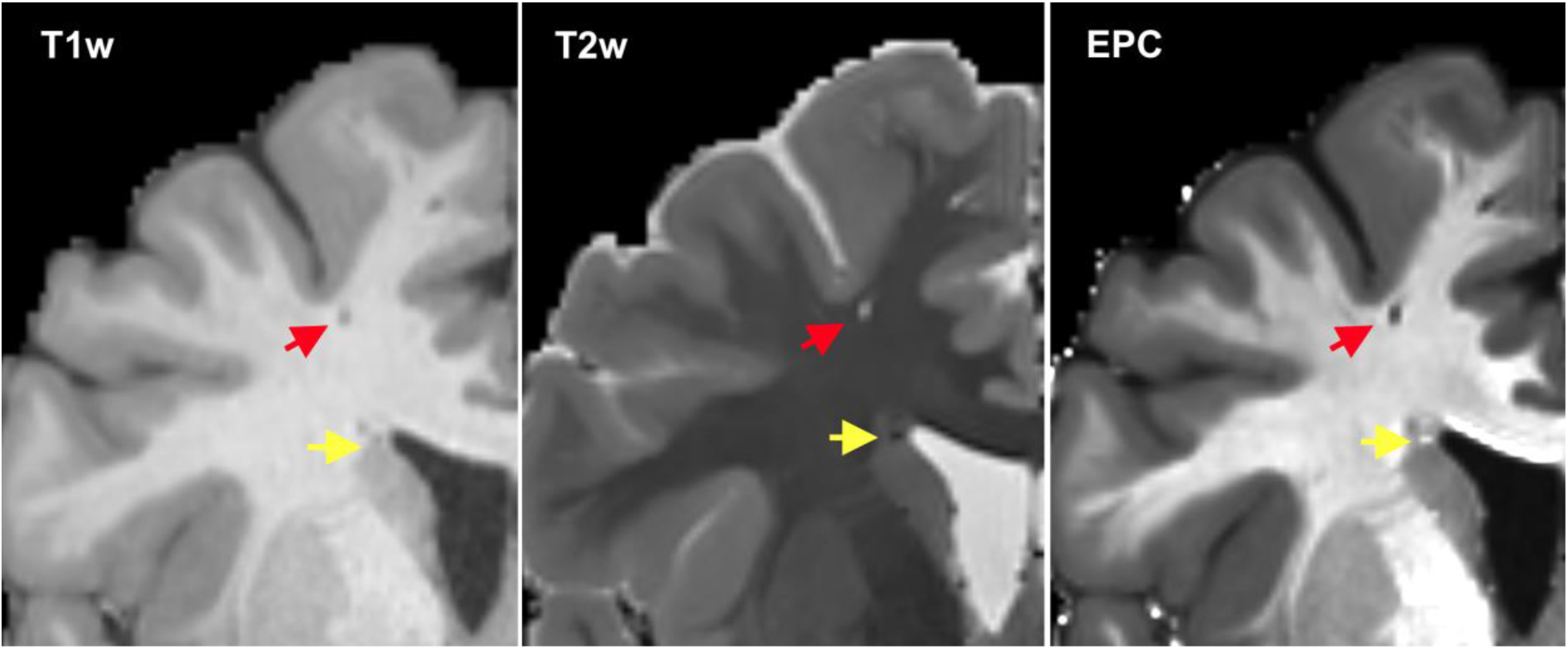
Different signature of vessels with and without PVS on the EPC. Red arrow shows a vessel with PVS. Yellow arrow shows a vessel with reasonably no surrounding PVS, due to the absence of the PVS-like signal on T2w (i.e. bright signal). The bright and dark voxels indicated by the arrows were followed through multiple slices to ensure they are indeed vessels. Note that differentiation of the PVS would have been incorrect in an automated technique if only T1w was used.

